# Downregulation of Perilipin1 by IMD leads to LD reconfiguration and adaptation to bacterial infection in *Drosophila*

**DOI:** 10.1101/2020.04.30.070292

**Authors:** Lei Wang, Jiaxin Lin, Junjing Yu, Kaiyan Yang, Li Sun, Hong Tang, Lei Pan

**Affiliations:** Wuhan Institute of Virology, Chinese Academy of Sciences, Wuhan, Hubei 430071, China; The Center for Microbes, Development and Health; Key Laboratory of Molecular Virology and Immunology, Institut Pasteur of Shanghai, Chinese Academy of Sciences, Shanghai 200031, China; CAS Center for Excellence in Biotic Interactions, University of Chinese Academy of Sciences, Beijing 100049, China; Shanghai Institute of Biochemistry and Cell Biology, Chinese Academy of Sciences, Shanghai 200031, China; University of Chinese Academy of Sciences, Beijing 100049, China

## Abstract

Lipid droplets (LDs) are dynamic intracellular organelles critical for lipid metabolism. Dynamic alterations in the configurations and functions of LDs during innate immune response to bacterial infections and the underlying mechanisms however, remain largely unknown. Herein, we trace the time-course morphology of LDs in fat bodies of *Drosophila* after transient bacterial infection. Detailed analysis shows that *perilipin1* (*plin1*), a core gene regulating lipid metabolism of LDs is suppressed by IMD/Relish, an innate immune signaling. During immune activation, downregulated *plin1* promotes the enlargement of LDs, which in turn alleviates immune reaction-associated reactive oxygen species (ROS) stress. Thus, the growth of LDs is likely an active adaptation to maintain redox homeostasis in response to IMD activation. Therefore, our study provides evidence that *plin1* serves as a modulator on LDs’ reconfiguration in regulating infection-induced pathogenesis, and Plin1 might be a potential therapeutic target for coordinating inflammation resolution and lipid metabolism.

## INTRODUCTION

Immune activation is essentially accompanied by metabolic reprogramming, which redistributes accessible energy to prioritize immune protection against pathogenic infections (1, 2). Thus, stringent regulation of metabolic machinery in response to immunoreaction is critical for the host fitness. Besides carbohydrates, lipids provide another important bioenergetic and synthetic resource to the host. Not limit to this, a number of lipid metabolites in turn have been reported to play key roles in pro- or anti-inflammatory pathways (3–5). In all eukaryotic and some prokaryotic cells, there are important intracellular organelles, lipid droplets (LDs), which provide a major place for the synthesis, lysis, transfer and storage of lipids or their derived metabolites (6). LDs contain a hydrophobic core of neutral lipids, such as di/triacylglycerols or sterol esters, which is surrounded with a phospholipid monolayer decorated by different proteins (7). Previously, LDs were considered to get involved in many physiological and pathological processes just because of their main functions on storing/providing energy and/or buffering toxic lipid species through modulating enzymic or autophagic lipolysis (8, 9). So far, emerging evidences have shown that LDs also take part in immune regulation. For instance, LDs modulate functions of myeloid cell through immune-metabolic reprogramming (10). LDs facilitate hosts to combat pathogens’ infections through selectively recruiting immune proteins (11, 12). Recently, a study indicated that mammalian LDs respond to bacterial lipopolysaccharide and function as innate immune hubs to coordinate host defense and cell metabolism (13). However, the role of LDs as pro- or anti-inflammatory modulators is still controversial (14), due to fact that LDs are highly dynamic organelles. The number, size and anchored proteins of LDs change quickly in response to infection or stress (15–17). These evidences also indicate that the status of LDs should be tightly controlled. Defects in the biogenesis and mobilization of LDs not only result in lipotoxicity (18, 19) but also exacerbate inflammatory responses and organelles dysfunction (20, 21). However, the role and dynamic pattern of LDs during immune process is still barely described. Especially, the factors mediating the transformation of LDs underlying immunometabolic switches have not been well identified.

LDs are non-homogenous organelles, which accommodates hundreds of variable proteins (22, 23). However, Perilipins (Plins) are the most prominent proteins that span the surface of LDs (24, 25). Each LD is usually decorated by two or more members of Perilipin family proteins and no LDs without perilipins have been identified in mammalian cells so far (26). There are five major perilipins in mammalian, named Perilipin1-5. These Perilipins differ in the expression and cellular localization in different tissues and have essential roles in the regulation of LDs’ structure and morphology (27, 28). However, whether Perilipins get involved in immune functions, probably through mediating LDs’ reconfiguration, is still obscure. In human or mouse adipocyte tissue, Perilipin1 deficiency leads to uncontrollable LDs lipolysis and infiltration of inflammatory cells (29, 30). Inhibition of lipases, such as adipose triglyceride lipase (ATGL) or hormone-sensitive lipase (HSL), can alleviate this metaflammation (31, 32). *Plin1* knockout also promotes secretion of prostaglandins, the pro-inflammatory lipid metabolites, and elevates pro-inflammatory M1-type adipose tissue microphages in mice (33). In innate immunity, the expression and localization of Perilipins on LDs changed in response to LPS stimulation, which subsequently affected antimicrobial capacity (13). These evidences suggest a link between Perilipins and immunometabolic regulations.

*Drosophila melanogaster* has emerged as a productive organism to investigate immunometabolism, due to the advantages of powerful genetic manipulation and highly conserved mechanisms in both innate immunity and metabolism (34–36). Especially, the fat body (analogous to human liver and adipose tissue), as a major organ mediating systemic innate immunity, is an ideal place for studying the interaction between metabolism and inflammation of LDs, due to its richness in LDs (37, 38). Furthermore, Perilipins are evolutionarily conserved from fungi to human. Not like more redundant Plins in mammalian, there are only two Plins in *Drosophila*, Lipid storage droplet-1 (*lsd1*or *plin1*) and *lsd2* (*plin2*) (39). Plin2 acts to promote lipid storage and LDs’ growth as a barrier for lipase (40–42), while Plin1 modulates protein flux on LDs (27, 28). They have opposite functions on the control of LDs’ morphology (27). In *Drosophila*, the immune deficiency (IMD) pathway is a dominant innate immune singling against Gram-negative bacterial infections, which is homologue to mammalian NF-kB/TNF pathway (35). Couple studies have revealed that IMD signaling modulate lipolysis in either fat body or intestine of flies (43, 44). And thus, LDs’ accumulation was once reported in fly gut after IMD activation (17). However, as if LDs adapt to immune response, whether and how morphological changes occur on LDs remain unclear. More importantly, the contribution of theses adaptive altered LDs to infectious pathogenesis and the underlying mechanisms mediating by LDs-anchored factors such as Plins, are still poorly understood.

In this study, the alteration in morphology and number of LDs were traced dynamically during bacterial infection. Plin1 was found to respond to IMD action and then modify LDs’ morphology to alleviate inflammatory stress. Our data reveal that adaptive modification of LDs acts as an active modulator of infection-induced pathogenesis.

## Results

### Bacterial infection modulates lipid metabolism, and particularly alters morphology of lipid droplets (LDs) in the fat body

In *Drosophila*, the fat body is not only a central organ mediating systemic immune responses, but also the epicenter for lipid metabolism. Thus, to decipher the mechanistic connections between innate immunity and lipid metabolism, the kinetics of fat content was tested in the fat body of *Drosophila* after systemic infection. *Escherichia. Coli* (*E. coli*), a non-pathogenic Gram-negative bacterium to flies, was used to perform nano-injection to infect adult male fruit flies. The immune deficiency (IMD) pathway is a dominant innate immune signaling against Gram-negative bacterial infections that regulates Relish/NF-κB-dependent transcription of AMPs, such as *Diptericin* (*Dpt*) (35). Thus, by measuring the expression level of *Dpt,* IMD signaling activity could be monitored (45, 46). In consistent with previous report that *E.coli* injection resulted in a transient innate immune response within 48 hour post infection (hpi)(47), a gradient increase in IMD activity in the fat body was observed from 0 hpi to 12 hpi, and then this activity subsided to the basal level after 48 hpi (**Fig.1A**). Interestingly, compared to mock injection control (**Supplementary Fig. S1A**), the fat levels in the fat body of flies with *E.coli* infection steadily increased from 4 hpi to 16 hpi and then almost recovered after 48 hpi (**Fig. 1A**). Therefore, these results suggest a link between lipid metabolism and IMD signaling activation in the fat body of flies.

**Fig. 1.**
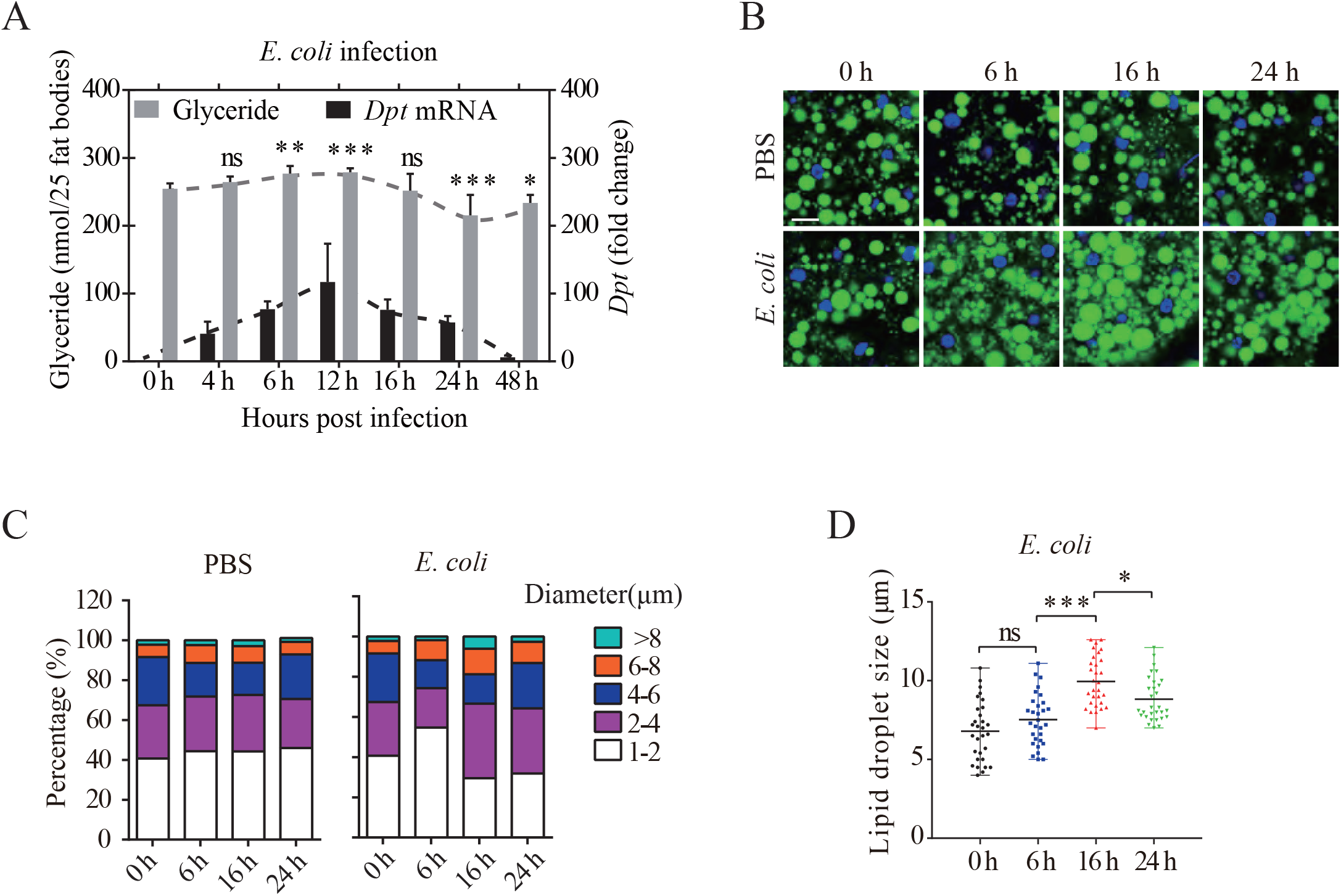
*E. coli* infection switches lipid metabolism and LDs morphology in the fat body. **(A)** Relative *Dptericin*(*Dpt*) mRNA expression and glyceride level in the fat body of wild type flies at indicated time points post *E. coli* infection. The mean values of *Dpt* mRNA expression or glyceride level was connected by dash line. The fold change of mRNA expression was normalized to that of 0 h and four independent repeats (n =20 flies per repeat) were performed at each time point. Total glyceride level of 25 flies’ fat body tissues was quantified in six biological replicates at each time point. (**B and C**) BODIPY staining (green) of LDs in the fat body of wild type flies at indicated time points post *E. coli* infection. Nuclei of fat body cells were stained with DAPI (blue). Scale bar: 10 μm. The corresponding statistics of the distribution of LDs’ size was shown in (**C**) for *E. coli* infection (n =30 cells for each time point). Eight fat bodies were examined for each time point.**(D)** The statistics of LDs’ size (n =30 cells) in the fat body of wild type flies at indicated time point post *E. coli* infection. Each scattering dot represents the data from one fat body cell. Error bars represent the mean ± s.d. (**A-**) and mean with range (**D**). Data were analyzed by One-way ANOVA with Tukey’s multiple-comparison test (**A, D**). \**p* < 0.05; \*\**p* < 0.01; \*\*\**p* < 0.001; ns, not significant. **See also in Supplementary Figure 1**.

LDs are the main site for lipid metabolism, mobilization and storage (48), which prompted us to investigate whether LDs change in the fat body in response to bacterial infection. BODIPY staining of fat body cells revealed that compared to PBS injection group (**Fig. 1B and 1C**), *E. coli* infection increased the percentage of intracellular small LDs (diameter < 2 μm) at 6 hpi (**Fig. 1B and 1C**). And then, LDs grew bigger at 16 hpi as indicated by the decrease in the percentage of small LDs and concurrent increase in the percentage of large LDs (diameter > 4 μm). Finally, this size distribution of LDs was restored to basal levels at 24 hpi (**Fig. 1B and 1C**). Accordingly, the average size of LDs in fat body cells had the similar changing trend (**Fig.1D**). These results indicate that small LDs are prone to fuse into bigger ones during the initial 16 h after *E. coli* infection.

### The reconfiguration of LDs requires IMD signaling activation

To determine whether IMD signaling activation rather than live bacterial growth is responsible for the modification of LDs during infection, heat-killed *E. coli* was applied to repeat infection in wild type flies. The elevated fat levels in fat bodies were still observed at 12 hpi in WT flies (**Fig. 2A**). In *Drosophila*, peptidoglycan (PGN) from Gram-negative bacteria can bind to the receptor of PGRP-LC to active IMD signaling through transcriptional regulator Relish (35). Thus, the flies with homozygous mutation of *PGRP-LC* (*PGRP-LC*^*Δ5*^) or *relish* (*relish*^E20^) was used for infection. In contrast to wild type flies, the phenotype of elevated fat contents in fat bodies disappeared and even reversed in these mutant flies at 12h post heat-killed *E.coli* injection (**Fig. 2B**). Moreover, IMD signaling deficiency also restricted the increase in LDs size at 16 hpi, compared to WT controls (**Fig. 2C and D**). Therefore, these results suggest that IMD signaling activation is required to modify LDs’ morphology in response to bacterial infection.

**Fig. 2.**
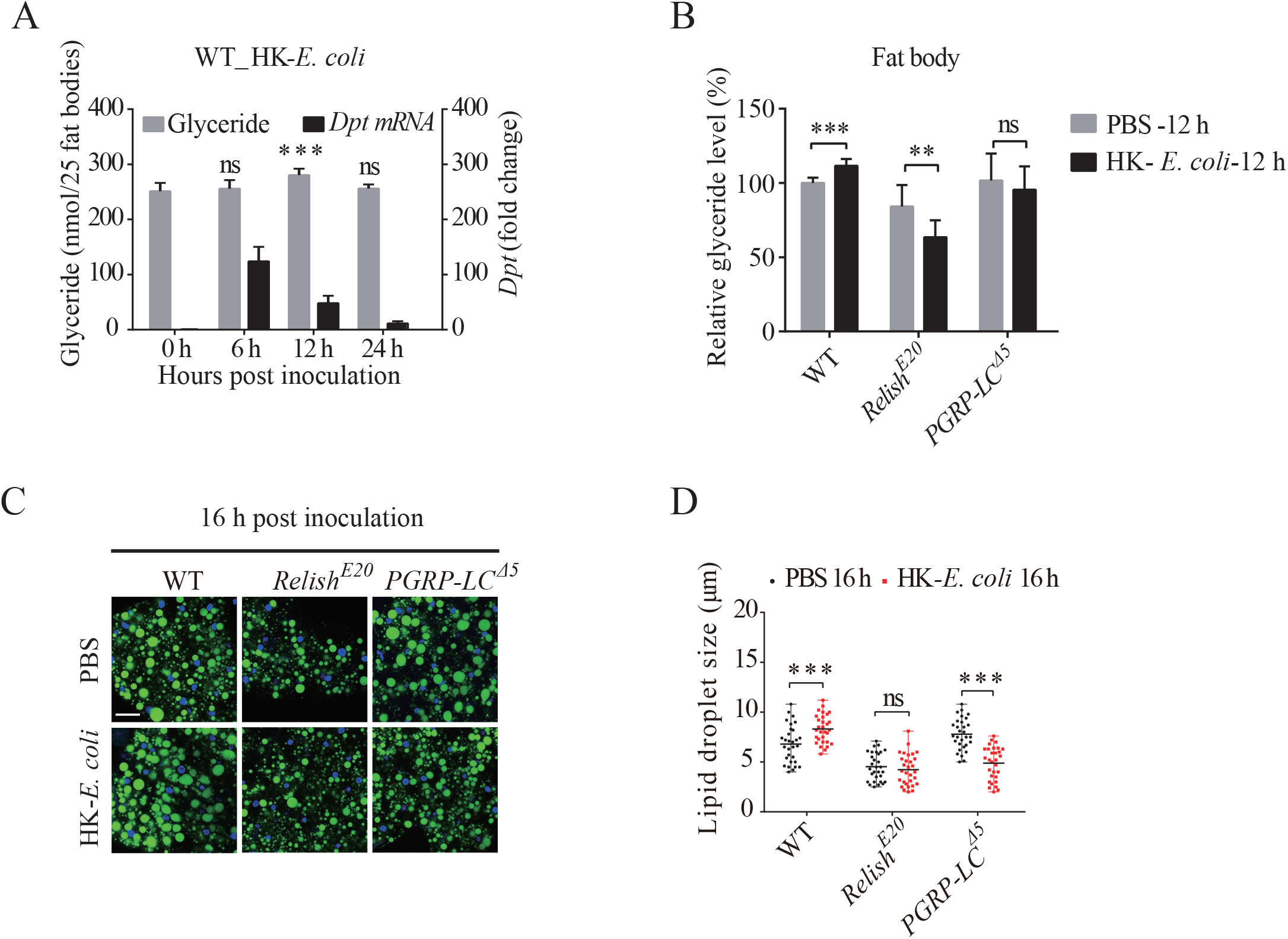
IMD signaling pathway is required for the alteration of lipid metabolism and LDs morphology upon infection. (**A**) Relative *Dptericin* (*Dpt*) mRNA expression (black) and Fat levels (Grey) in the fat body of wild type flies at indicated time points post heat-killed *E. coli* (HK-*E. coli*) infection. The fold change of mRNA expression was normalized to that of 0 h and four independent repeats (n =20 flies per repeat) were performed at each time point. Total fat levels of 25 flies’ fat body tissues was quantified in six biological replicates at each time point. (**B**) Relative glyceride level in the fat body of wild type and IMD pathway mutants (*Relish* and *PGRP-LC*). Each value of glyceride level was normalized to that of 0h of wild type. Each data contains four independent repeats (25 flies’ fat body tissues per repeat). (**C** and **D**) BODIPY staining (green) of LDs (**C**) and the corresponding statistics of LDs’ size (n =30 cells) (**D**) in the fat body of IMD pathway mutant flies and corresponding genetic control flies. Eight fat bodies were examined for each sample. Error bars represent mean ± s.d. (**A-B**) or mean with range (**D**). Data were analyzed by One-way ANOVA with Tukey’s multiple-comparison test (A-B) and Multiple t-tests (D). Scale bar: 20 μm. \**p* < 0.05; \*\**p* < 0.01; \*\*\**p* < 0.001; ns, no significance.

### *plin1* is involved in LDs’ morphological change induced by IMD activation

Perilipins (Plins), a group of constitutive proteins that span the surface of LDs, were reported to regulate lipid mobilization and LDs’ morphology (27, 28). There are two Perilipins in *Drosophila*, Plin1 and Plin2. To explore whether Plins are involved in the regulation of LDs’ reconfiguration in response to immune activation, their time-course expression was detected in the fat body by real-time PCR. *E. coli* infection induced a significant downregulation of *plin1* mRNA levels at 4 hpi, which was then gradually restored to basal levels at 24 hpi (**Fig.3A**). The changing trend of *plin1* expression seemed to be negatively correlated with the changes in LD’s size and IMD activity. However, the expression level of *plin2* was only slightly tuned down at 4hpi and back to normal at 12h after *E. coli* infection (**Supplementary Fig.2A**). Previous study have shown that deficiency of *plin2* resulted in reduced rather than enlarged size of LDs (27). Therefore, these results indicate a potential role of *plin1* in the regulation of LDs’ morphology in response to IMD activation. Furthermore, either deficiency of *plin1* by mutation (*plin1*^38^) or specific knockdown of *plin1* in fat body (*UAS-plin1 RNAi* driven by *ppl-GAL4*) promoted the formation of large LDs. Whereas, ectopic expression of *plin1* in the fat body led to the accumulation of much smaller LDs, compared with controls (**Fig. 3B**). These results were reminiscent of previous studies that Plin1 may function to enhance lipid mobilization and inhibit LD coalescence (49).

**Fig. 3.**
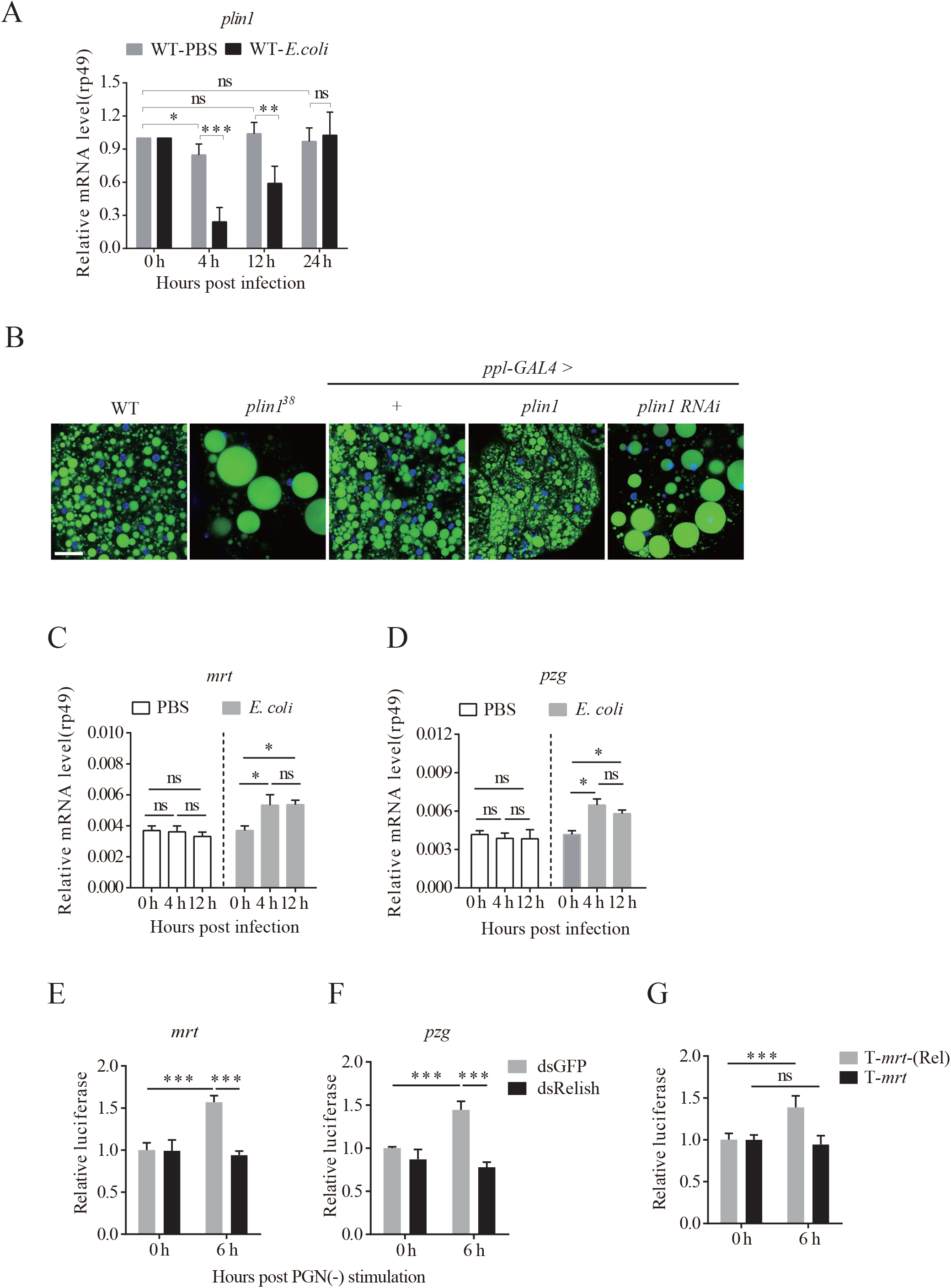
*plin1* responds to IMD activation through Mrt/Pzg complex, and regulates LDs’ morphology. (**A**) Relative *plin1* mRNA levels in the fat body of wild type flies at the indicated time points post *E. coli* infection. Flies treated with sterile PBS were used as a control. The fold change of mRNA expression was normalized to that of 0 h. (**B**) BODIPY staining (green) of LDs in the fat body of indicated flies. Nuclei of fat body cells were stained with DAPI (blue). Eight fat bodies were examined for each genotype. (**C** and **D**) Relative *mrt* (**C**) and *pzg* (**D**) mRNA levels in the fat body of wild type flies post *E. coli* infection. Flies treated with sterile PBS were used as a control. The fold change of mRNA expression was normalized to that of 0 h. Four independent repeats (n=20 flies fat body tissues per repeat) were performed at each time point for each group. (**E** and **F**) Relative luciferase activities of *mrt* (**E**) (Full length promoter of −1.5kb to +1bp including all predicted Relish Binding motifs in Fig. S3) reporter or *pzg* (**F**) (1.5 KB upstream of TSS, all predicted Relish binding sites are covered) reporter in S2* cells after double strand RNA (dsRNA) and PGN (35 μg/ml) treatment. All data were normalized to dsGFP control group at 0 h. (**G**) Relative luciferase activities of T-*mrt* (Rel) and T-*mrt* reporter in S2* cells after PGN (35 μg/ml) treatment. All data were normalized to T-*mrt* (Rel) group at 0h. Three independent repeats were performed at each time point for each treatment. Error bars represent the mean ± s.d.. Data were analyzed by One-way ANOVA with Tukey’s multiple-comparison test (**A, C-D**), Multiple t-tests (**A**) and Student’s t test (**E-G**). Scale bar: 20 μm. \**p* < 0.05; \*\**p* < 0.01; \*\*\**p* < 0.001; ns, no significance. **See also in Supplementary Figure 2 and 3**.

Martik (MRT) /Putzig (PZG) complex, a chromosome remodeling complex, has been reported to suppress *plin1* at transcriptional level (49). The mRNA levels of both *mrt* and *pzg* were upregulated in the fat body after bacterial infection (**Fig. 3C and 3D**). Interestingly, homologous alignment showed that at least one conserved binding motif of Relish existed in the promoter region of both *mrt* and *pzg* genes across *Drosophila* species with different evolutionary ages (**Supplementary Fig. S3A and S3B**). This implies a potential regulation of these genes by IMD/Relish. Peptidoglycan (PGN) derived from gram-negative bacteria can activate IMD signaling in *Drosophila* S2* cells *in vitro* (50). The treatment of PGN enhanced luciferase activity controlled by the promoter of *mrt* or *pzg* in S2* cells, which was blocked by the knockdown of Relish using dsRNA (51) (**Fig. 3E and 3F**). Additionally, two Relish binding motifs in truncated *mrt* promoter region (T-*mrt(Rel)*, −870 to +1bp, in **Supplementary Fig. S3C**) were required for *mrt* transcription (**Fig. 2G**), because PGN treatment didn’t enhance T-*mrt*-Luc activity any more when these two sites were removed (**Fig. 2G**). Thus, these results suggest that suppression of *plin1* by IMD signaling might be through upregulation of mrt/pzg. All together, these results provide an explanation for LDs’ growth in the early stages of transient IMD activation.

### Plin1 compromises host protection against bacterial infection

Naturally, whether Plin1 participated in the defense of bacterial infection was tested next. Since *E. coli* is non-pathogenic to flies, another Gram-negative bacterium, *Salmonella. Typhimurium* (*S. typhimurium*), which is a deadly pathogen for flies (52), was used to evaluate Plin1 function on immune defense. Compared to genetic controls, either *plin1* deficiency (*plin1*^38^) (**Fig. 4A and 4B**) or fat body-specific knockdown of *plin1* (*ppl-GAL4*>*UAS-plni1RNAi*) (**Fig. 4C and 4D**) significantly prolonged the survival rate and slightly reduced bacterial loads (colony-forming units, CFUs) after *S. typhimurium* septic infection, indicative of enhanced resistance against bacterial infection. Conversely, ectopic expression of *plin1* in the fat body (*ppl-GAL4*>*UAS-plin1*) led to a dramatic increase in mortality rate of flies infected with *S. typhimurium* (**Fig. 4E**, Reducing infection OD because O.E.*plin1* flies died too quickly.), or even by non-pathogenic *E. coli* (**Fig. 4G**), possibly due to uncontrolled bacterial growth (**Fig. 4F and 4H**). However, it’s worthy to note that deficiency of *plin1* did not affect anti-microbial peptides (AMPs) (*Diptericin*, *Dpt; AttacinA, AttA*) response upon *E. coli* infection (**Supplementary Fig. S4A**), but specifically improved *Dpt* expression upon *S. typhimurium* infection (**Supplementary Fig. S4B**). Interestingly, overexpression of *plin1* dampened AMPs response in both *E. coli* and *S. typhimurium* infections (**Supplementary Fig. S4C and S4D**). Taken together, these results suggest that adaptive downregulation of *plin1* in response to IMD signaling activation protected the host against bacterial infections.

**Fig. 4.**
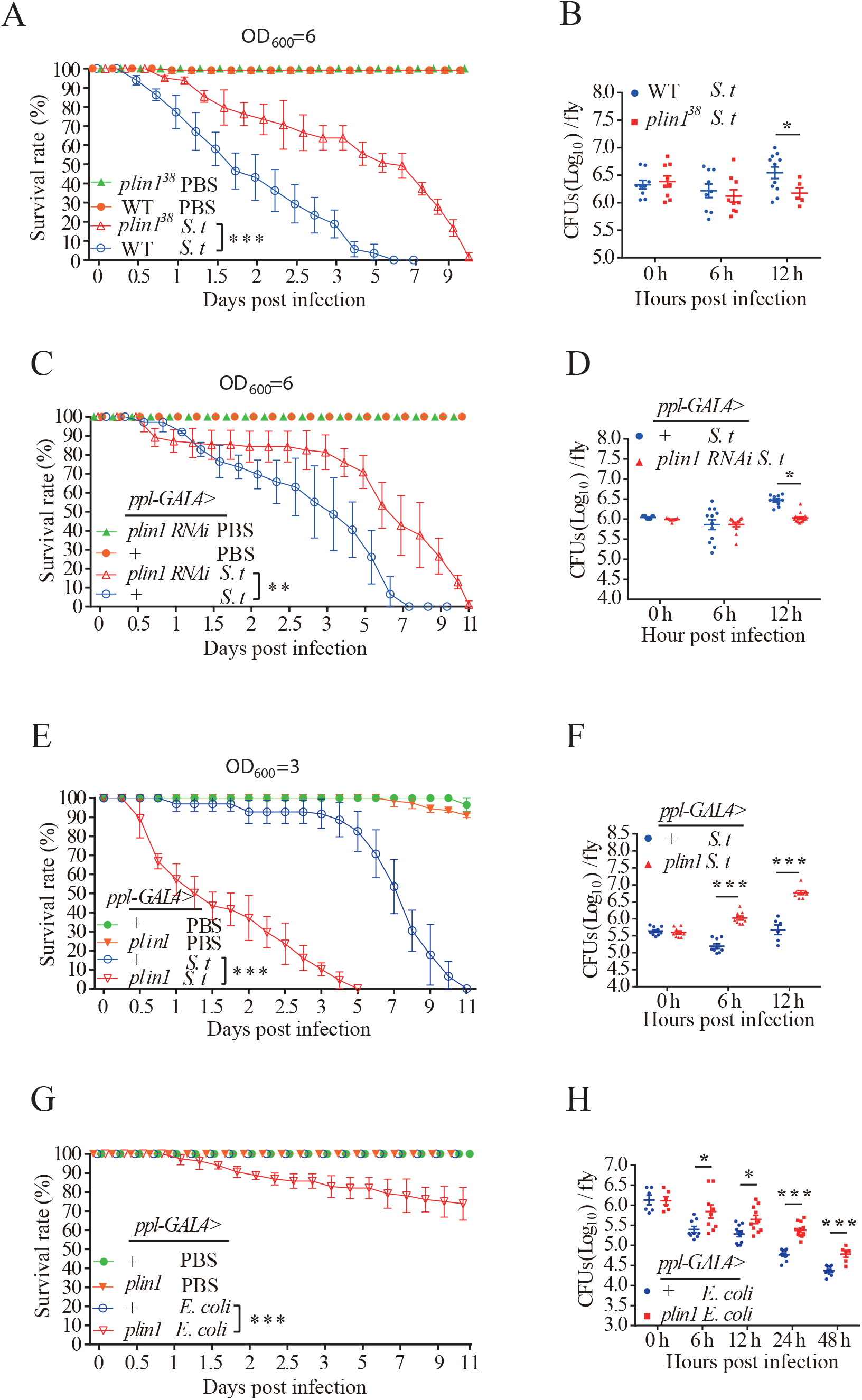
*plin1* participates in the susceptibility of flies to bacterial infection. (**A** and **B**) Survival curves (**A**) and bacterial loads (CFUs) (**B**) of wild type and *plin1*^38^ flies (n =60) post *S. typhimurium* infection. (**C** and **D**) Survival curves (**C**) and bacterial loads (CFUs) (**D**) of *ppl-GAL4>plin1 RNAi* and control flies (n =60) post *S. typhimurium* infection. (**E** and **F**) Survival curves (**E**) and bacterial loads (CFUs) (**F**) of *ppl-GAL4>plin1* and control flies (n =60) post *S. typhimurium* infection. (**G** and **H**) Survival curves (**G**) and bacterial loads (CFUs) (**H**) of *ppl-GAL4>plin1* and control flies (n =60) post *E. coli* infection. Values of plotted curves represent mean ± s.d. (**A, C, E, G**) of at least three independent repeats. Each scattering dot (CFUs) represents one technical replicate, line represents the mean of four independent repeats (**B, D, F, H**). Data were analyzed by Kaplan–Meier (**A, C, E, G**) and Multiple t-tests (**B, D, F, H**). \**p* < 0.05; \*\**p* < 0.01; \*\*\**p* < 0.001, ns, no significance. **See also in Supplementary Figure 4**.

### Plin1-mediated reconfiguration of LDs participates in the homeostasis of intracellular ROS

The next question is whether downregulated Plin1-induced LDs’ growth also benefit the host against bacterial infection. Sustained immune activation is a high energy-cost process, which requires active lipolysis and usually leads to excessive reactive oxygen species (ROS) accumulation due to the release and oxidation of free fatty acids, one hallmark for inflammatory damages (18, 53). However, LDs’ growth could efficiently reduce the accumulation of free fatty acids, and probably relieve ROS-related tissue damages (54). As expect, *plinl1* deficiency (*plin*^138^) (**Fig.5A and A1**) or knockdown (*UAS-plin1RNAi* driven by *ppl-GAL4*)(**Fig. 5B and B1**), which promoted LDs growth, accompanied with a much lower level of ROS than that of control. In contrast, overexpression of *plin1* in fat body cells(*ppl-GAL4*> *plin1*), which transformed LDs into smaller ones, markedly increased ROS intensity (**Fig. 5B and B1**). These results suggest a correlation between the size of LDs and the intensity of ROS accumulation in fat body cells. To further support this notion, we skewed ROS metabolism in fat bodies through knockdown of <u>s</u>uper<u>o</u>xide <u>d</u>ismutase genes (*sod1* or *sod2*) or <u>cat</u>alase gene (*cat*), all of which encode enzymes for intracellular ROS clearance (55, 56). All these flies (*ppl-GAL4>UAS-sod1-RNAi*, *sod2-RNAi* or *cat-RNAi*) contained elevated ROS levels (**Fig.5C and 5D**) and compensatory LD’s growth in fat bodies (**Fig. 5E and 5F**). If blocking the large LDs’ formation by simultaneous overexpression of *plin1* in these genetic backgrounds (**Fig. 5E and 5F**), much higher ROS accumulation was observed in fat bodies (**Fig. 5C and 5D**). All together, these results suggest that Plin1-controled LDs’ reconfiguration takes part in antioxidative functions.

**Fig. 5.**
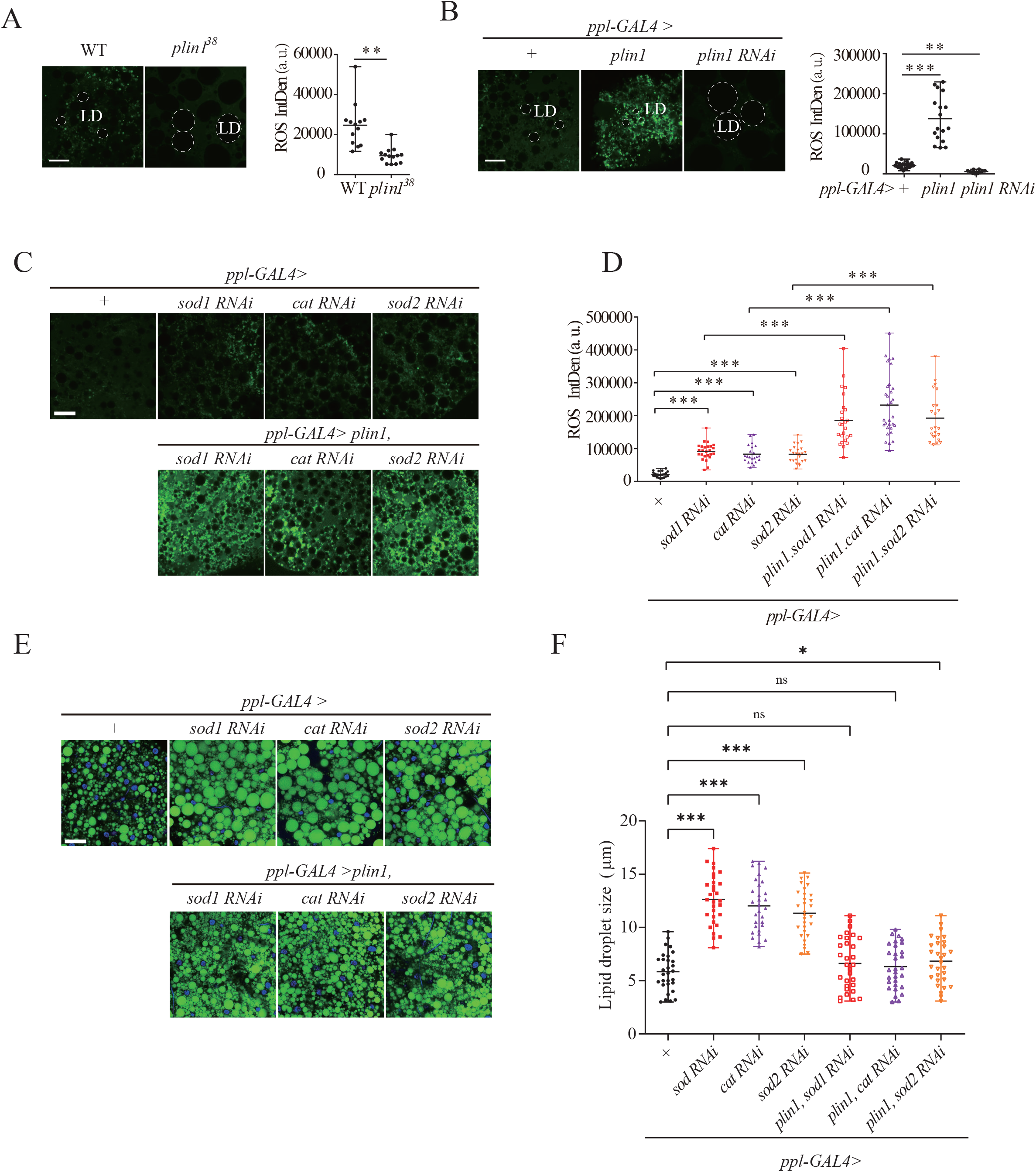
Plin1-mediated reconfiguration of LDs is involved in the regulation of intracellular ROS. (**A** and **B**) ROS level indicated by DCFH-DA staining (green) in the fat body of indicated flies. The statistics of fluorescence intensity was plotted in (**A**) for *plin1*^38^ and in (**B**) for *ppl-GAL4>plin1* and *ppl-GAL4>plin1-RNAi* flies and their genetic controls, respectively. Dashed circle indicated LDs. Eight fat bodies were examined for each sample and each scattering dot represents the data from one image. (**C** and **D**) ROS levels indicated by DCFH-DA staining (green) in the fat body of indicated one-week old adult flies and control flies (**C**). The corresponding fluorescence intensity was quantified in (**D**). Eight fat bodies were examined for each sample and each scattering dot represents the data from one image. (**E** and **F**) BODIPY staining (green) of LDs (**E**) and the corresponding statistics of LDs’ size (n =30 cells) (**F**) in the fat body of indicated flies. Eight fat bodies were examined for each sample. Error bars represent the mean with range. Data was analyzed by Student’s t test (**A-B, D, F**) and One-way ANOVA with Tukey’s multiple comparison test (**D, F**). Scale bar: 20 μm. \**p* < 0.05; \*\**p* < 0.01; \*\*\**p* < 0.001, ns, no significance.

### Downregulated Plin1 in response to IMD activation benefits flies against oxidative stress associated with bacterial infection

The activation of immune signaling, such as NF-κB or TNF signaling, often associates with ROS-induced inflammatory stress (53, 57). A transgenic allele with a *gstD-GFP* insertion was utilized to monitor ROS activity *in vivo* by measuring GFP intensity (58). Indeed, in *Drosophila*, infection either by non-pathogenic *E.coli* or by strong pathogenic *S. typhimurium* induced an obvious increase of intracellular ROS levels in the fat body (**Fig. 6A and 6B**). These results support the link between bacterial infection and accumulation of intracellular oxidative stress. If we removed this infection-associated intracellular ROS by feeding flies with N-acetylcysteine (NAC), a widely-used ROS scavenger, the survival of flies after pathogenic *S. typhimurium* infection was improved, compared to non-infected controls (**Fig. 6C**). It’s worthy to note that feeding flies with NAC at 12 h, not 0h, post *S. typhimurium* infection, benefited the fitness of flies much better. It’s likely that at time point of 12 hpi, excessive ROS accumulation had already developed, and initiate ROS of early infectious stage is useful for defense against bacteria (59, 60). These results suggest that excessive oxidative stress, which develops during bacterial infection is harmful for the host.

**Fig. 6.**
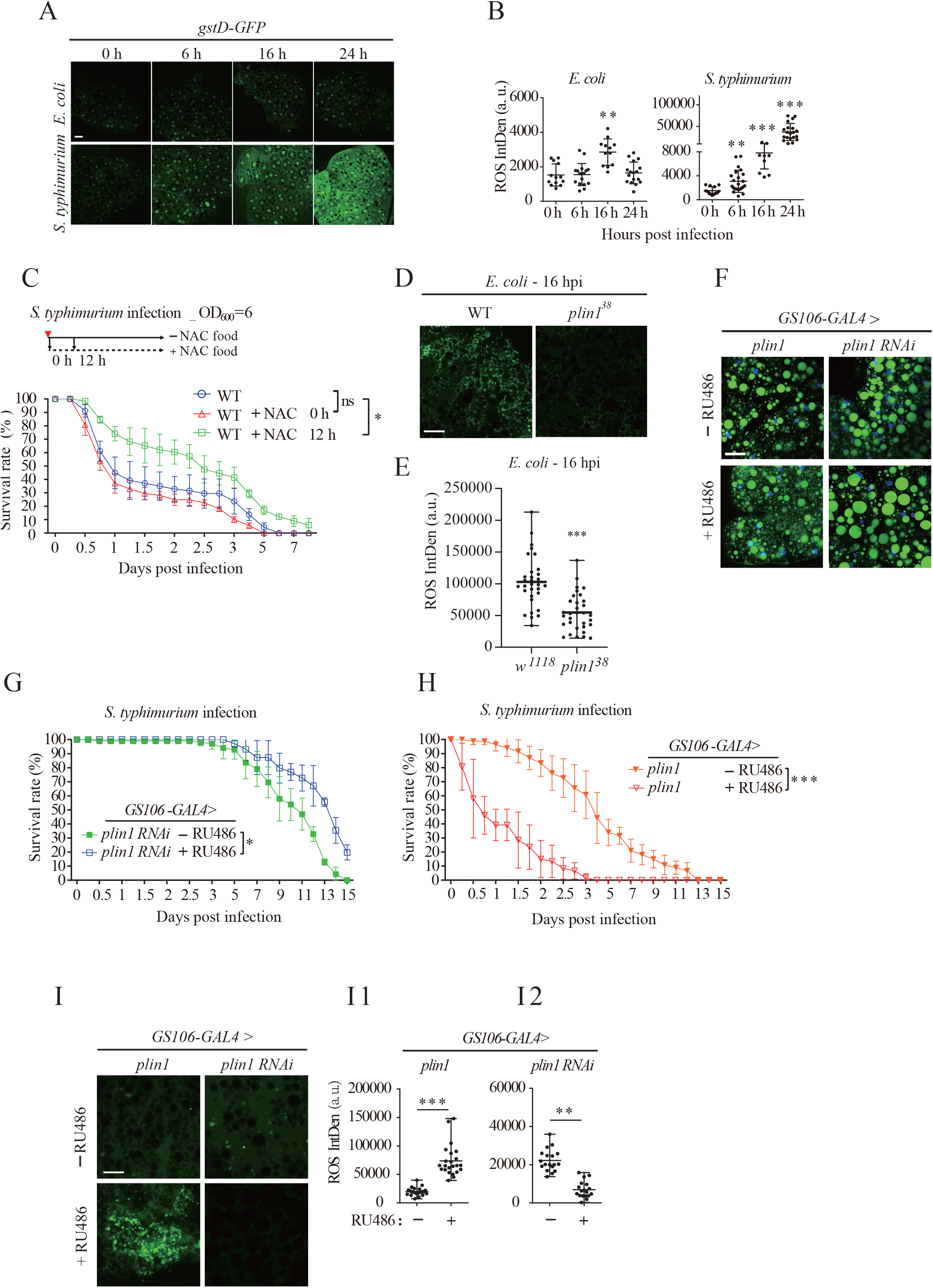
Downregulated Plin1 benefits flies against bacterial infection through reducing oxidative stress. (**A** and **B**) ROS level indicated by GFP intensity (green) of *gstD-GFP* reporter in the fat body of wild type flies infected with *E. coli* (upper panel) or *S. typhimurium* (lower panel) at indicated time points. The statistics of GFP intensity was plotted in (**B**) for *E. coli* infection and *S. typhimurium* infection. Eight fat bodies were examined for each sample and each scattering dot represents the data from one image. (**C**) Survival curves of wild type flies (n=60 flies) with or without NAC treatment at indicated time post *S. typhimurium* infection. (**D** and **E**) ROS level indicated by DCFH-DA staining (green) in the fat body of indicated flies at 16 h post *E.coli* infection. (**F**) BODIPY staining (green) of LDs in the fat body of *GS106-GAL4>plin1* and *GS106-GAL4>plin1 RNAi* flies after treatment with (lower panel) or without (upper panel) RU486 treatment. Eight fat bodies were examined for each sample. (**G** and **H**) Survival curves of above flies (n =60) post *S. typhimurium* infection. (**I**) ROS level indicated by DCFH-DA staining (green) in the fat body of indicated flies with (lower panel) or without (upper panel) RU486 treatment. The statistics of fluorescence intensity was plotted in (**I1**) for *GS106-GAL4>plin1* and in (**I2**) for *GS106-GAL4>plin1 RNAi* flies. Eight fat bodies were examined for each sample and each scattering dot represents the data from one image. Error bars represent the mean with range (**B, E, I**). Values of plotted curves represent mean ± s.d. of at least three independent repeats (**C, G-H**). Data was analyzed by One-Way ANOVA with Tukey’s multiple-comparison test (**B**), Student’s t test (**E, I**) and Kaplan–Meier (**C, G-H**). Scale bar: 20 μm. \**p* < 0.05; \*\**p* < 0.01; \*\*\**p* < 0.001, ns, no significance.

Since LDs are major hubs for lipid metabolism in fat body cells and function on ROS clearance, it promoted us to investigate whether *plin1*-mediated LDs modification is involved against bacterial infection, probably through regulating intracellular ROS. At first, we found that elevated ROS levels in wild type fat bodies were diminished in fat bodies of *plin1* deficiency flies (**Fig. 6D and 6E**). Next, a RU486 induced fat body-specific GAL4 (*GS106-GAL4*) was used to modulate the expression of *plin1* just before *S.typhimurium* infection, which exclude the possible effects of *plin1* on the development of flies. As expect, overexpression or knockdown of *plin1* in the fat body resulted in a significant decrease or increase in LDs’ size, respectively (**Fig. 6F**). Downregulation of *plin1* prolonged the survival rate after *S.typhimurium* infection (**Fig. 6G**), while ectopic expression of *plin1* shortened the life span dramatically (**Fig. 6H**). Meanwhile, fluorescent probe 2’,7’-dichlorofluorescein diacetate (DCFH-DA) staining indicated an elevated intracellular ROS levels in *plin1*-overexpression flies and a reduced ROS levels in *plin1*-knockdown flies during infection (**Fig. 6I and 6I1-2**). Taken together, these results suggest that large LDs formation contributes to alleviate intracellular oxidative stress induced by bacterial infection and Plin1 might serve as an important modulator to promote LDs modification in response to IMD activation.

## Discussion

Metabolic reprogramming of lipids has been widely reported to be associated with immune responses (1, 2, 61). As a major intracellular organelle for lipid metabolism and storage, LDs also seem to be involved in immune processes. Immune stimulation either by infection with bacteria (62, 63), virus (64–66), fungus(67) or protozoan parasites (68), or by cytokines inoculation (69, 70) may promote the biogenesis of LDs in mammalian leukocytes. Recently, Hash et al also reported that LDs are infection-inducible organelles in the gut of *Drosophila* at a certain timepoint after infection (17). Bosch et al further indicated LDs recruit antimicrobial proteins in response to LPS and function as innate immune hubs (13). However, the status and morphology of LDs change rapidly *in vivo*. Whether this dynamic transformation of LDs in response to immune stimulation is seldom described. Whether adaptive morphological change of LDs plays active rather than passive roles in pathogenesis, and key regulators linking LDs’ reconfiguration and infection still need further investigated.

In this study (**Fig.7**), we carefully traced the time-course morphogenesis of LDs in the fat body along with the dynamic curve of IMD signaling activity. We found that transient IMD activation by bacterial infection promoted LDs’ growth in the fat body within 12hpi. Both LDs’ size and fat levels in the fat body was maximum when IMD activity almost achieved its peak. Previous studies show that transcriptional levels of most triglyceride synthesis genes are suppressed during the initial phase of infection (71, 72), suggesting that the substrates for LDs’ biogenesis in the fat body were probably imported lipids rather than *de novo* synthesized fatty acids, and IMD signaling activation is required for this process. Detailed analysis showed that *plin1* downregulation is critical for LDs’ growth in response to transient IMD activation, considering its expression was suppressed by IMD/Relish activated MRT/PZG complex. Although the immune response is an energy-cost process, the fat content in specific tissue such as fat body is surprisingly increasing at the early stage of immune activation. These findings prompt us to imagine that LDs’ biogenesis is likely an active host adaptation to immune challenges. To further support this hypothesis, we found that enlarged LDs benefit the host against intracellular ROS-mediated oxidative stress induced by bacterial infection.

**Fig. 7.**
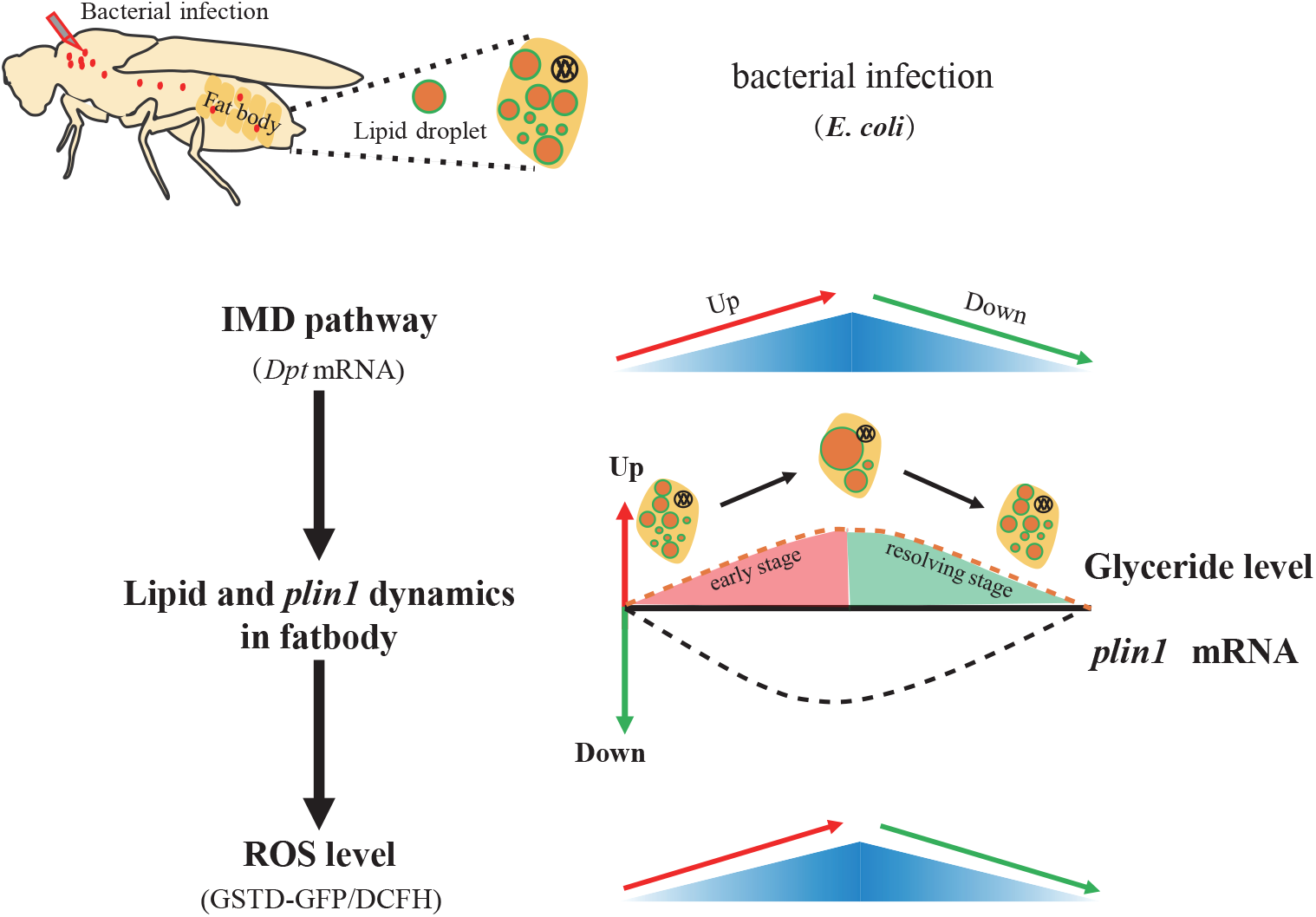
The schematic diagram of LDs’ morphogenesis mediated by Plin1 during infection-induced pathogenesis. LD’s growth induced by downregulation of *plin1* in response to IMD signaling activation post bacterial infection. And enlarged LDs provides antioxidant role and benefits the host for anti-infection.

Excessive ROS accumulation is often the main cause of inflammation/infection-induced cellular damages. In fact, biological processes such as cancer, neural activity, and inflammation are all energy-intensive, rely on robust fat metabolism, which releases large amounts of free fatty acids. The excess accumulation of free fatty acids in the cytoplasm promotes lipotoxicity and ROS-induced oxidative stress (73–75). The high levels of intracellular ROS can further promote lipolysis and free fatty acids release (76). This vicious circle finally drives the host to enter a severe metaflammatory state during chronic hyperinflammation, and consequently shorten lifespan. A recent study showed that renal purge of hemolymphatic lipids can efficiently prevent ROS-mediated tissue damage during inflammation (77). A similar antioxidant function of LDs was also reported in neuronal stem cell niche (78) and in cancer cells (79). In our study, blocking the breakdown and promoting the growth of LDs by downregulated *plin1* could efficiently eliminate ROS accumulation and prolong flies’ lifespan after bacterial infection. However, the detailed mechanisms how large LDs prefer to prevent ROS accumulation needs further investigation. One possibility is that the formation of large LDs sequesters the release of excessive free lipids, which oxidation contributes to the main source of ROS generation. Another possibility is that the larger the LDs are, the smaller the contact areas with mitochondria are. As mitochondria provides a major place for the oxidation of lipid usually supplied by LDs (9), reduced contacts between LDs and mitochondria might be another not-bad way to cut down ROS generation. Thus, LDs’ growth is beneficial for redox homeostasis of the host. The downregulation of Plin1 to promote the enlargement of LDs might be an effective host adaptation to resolve inflammation-associated stress in response to immune activation.

Moreover, large LDs’ formation can reduce the opportunity of pathogens to utilize free fatty acids for their own growth (80–82). This is possibly one reason why *plin1* deficient flies, owning bigger LDs, had lower bacterial loads after infection. In addition, larger LDs might contain more resident histones, a cationic protein, which has been reported to kill bacteria in a previous study (83). In mammals, IFN-γ treatment of *M. tuberculosis* infected bone-marrow derived macrophage (BMDM) can induce the formation of LDs, in which neutral lipids serve as a source to produce eicosanoids for enhancing host defense (84). A recent study made a detailed analysis that LDs recruit cathelicidin, a broad-spectrum antimicrobial peptide, in response to LPS stimulation (13). However, whether the change of LDs’ morphology alters the recruitment of LD-anchored proteins and underlying molecular and cellular mechanisms need further investigation. In our study, the reduced expression of genes encoded antimicrobial peptides was also found when *plin1* overexpression (**Supplementary Fig. S4C and S4D**) and *diptericin* was upregulated after *plin1* mutant flies infected with bacteria (**Supplementary Fig. S4B**). These results also suggest that a link between antimicrobial signaling with *plin1* directly or Plin1-mediated LDs’ modification indirectly. However, the large LDs’ formation in response to bacterial infection could in turn benefit the host to combat pathogens actively.

Plin1 is an important protein factor on the surface of LDs. It has been reported to control the mobilization of lipids on LDs’ surface by recruiting kinds of enzymes (85, 86) and sufficient to alter the morphology of LDs (27). In this study, we found that the expression of *plin1* rather than *plin2* preferred to be regulated by innate immune signaling. This provokes us to conceive that Plin1 may serve as a bridge to link immunity and lipid metabolism through modification of LDs. In response to transient immune activation, adaptative enlarged LDs benefit the host against pathogens and inflammation-induced stress. The alternation of the levels of mammalian PlINs protein on LDs’ surface was once mentioned after LPS stimulation (13). Our study provide a possibility that perilipins might response to immune signals and play an active role in infectious pathogenesis through transforming LDs. It is worthy in the future to trace and dissect the dynamic protein compositions on the surface of LDs along the different stages of inflammation, especially the proteins interact with Plin1. In summary, we found that the Plin1-mediated LDs’ morphological alteration is not only an adaptive consequence after bacterial infection, but also actively contributes to pathogenic regulation. Therefore, reconfiguration of LDs may provide a potential therapeutic target for resolution of inflammation.

## Materials and Methods

### Drosophila stocks and bacterial strain

All flies were propagated at 25°C on standard cornmeal food (1 L food contains 77.7 g cornmeal, 32.19 g yeast, 10.6 g agar, 0.726 g CaCl2, 31.62 g sucrose,63.2 g glucose, 2 g potassium sorbate and 15 ml 5% Tegosept),30%-55% humidity with a 12h/12h light/dark cycle. Fly resources that were used in this study as follows: w^1118^ were used as wild-type controls if no additional indication. *plin1*^38^, *UAS-plin1* mcherry, *UAS-plin1 RNAi* and *ppl-GAL4* were kindly gifted from Dr. Xun Huang (Institute of Genetics and Developmental Biology, CAS). *UAS-gstD-GFP* was kindly gifted from Dr. ZhiWei Liu (Shanghai Ocean University). *w*^*1118*^, *w[1118];P{w[+mW.hs]=Switch1}106*, *PGRP-L*^Δ5^,*Relish*^*E20*^ were obtained from Bloomington stock center. All flies used in this study were male. Two bacterial strain, *E. coli* (*DH5a*) and *S. typhimurium (SR-11)* (a gift from Dr. ZhiHua Liu, Institute of Biophysics, CAS) were used in this study.

### Cloning and double-strand RNAs

To construct the *mrt* and *pzg* reporter vector (*mrt*-luc and *pzg*-luc), the *mrt* and *pzg* promoter sequence (about −1500 bp or −1000 bp to 0 bp) was PCR amplified from Drosophila genomic DNA and introduced into *pGL3* vector (Progema) at HindIII restriction site by using recombination technology (Hieff Clone® Plus One Step Cloning Kit, YEASEN). All the plasmid constructs were verified by nucleotide sequencing. *pAC5.1*-renilla plasmid as a normalized reporter. Double-stranded RNAs (dsRNAs) against relish or GFP used in the luciferase reporter assay were synthesized using MEGAscript T7 kit (Invitrogen). Primers used for PCR amplification are listed in Supplementary Table 1.

### Infection and survival rate counting

Bacterial strains used in this study are *E. coli* (*DH5a*) and *Salmonella typhimurium* (*S. typhimurium*). Two days before infection, both bacteria from glycerol stocks were streaked onto Luria Broth (LB) agar plates and grown overnight at 37°C. The plate could be stored at 4 °C for up to 1 week. A single colony was inoculated to 6 ml fresh LB medium and grown at 37°C with shaking (200 rpm). Grow the bacteria to an OD600 of 0.7 to 0.8 (about 3.5 hours). The bacterial culture was pelleted with sterile phosphate-buffered saline (PBS) to the desired concentration. We injected 50.6 nl of bacterial suspension into dorsal prothorax of each fly with Nanoject II injector (Drummond). All flies used were 1 week old after eclosion. The final optical density (O.D. / ml) at 600 nm for injection were *E. coli* (O.D. 10) and *S. typhimurium* (O.D. 6 or O.D. 3). For *E.coli* infection, each fly obtained about 1×10^6 CFUs. For *S. typhimurium* infection, each fly obtained the lower dose (about 2×10^5 CFUs) or the higher dose (about 1×10^6 CFUs) according to the experiment design. Infected flies about 23 per vial were maintained at 25°C. Death was recorded at the indicated time point, and alive flies were transferred to fresh food every day for the survival analysis and CFUs assay.

### Bacterial loads assay

To monitor bacterial loads of the flies during infection, the number of colony forming units (CFUs) grown on LB agar plate was determined as follow: 5 living flies were randomly collected in a 1.5 ml EP tube, rinsed with 70% ethanol two times by vertex for 10s to sterile the surface adherent bacteria, then rinsed with sterile deionized water two times by vertex for 10 s, and then homogenized in 200 μl of sterile PBS with three fly body volumes of ceramic beads (diameter: 0.5 mm) in the Minilys apparatus (Bertin TECHNOLOGIES) at highest speed for 30 s. The suspensions obtained were then serially diluted in PBS and plated on LB agar. Specially noted for *S. typhimurium* plating, PBS was substituted with PBS + 1% Triton X-100. For the bacterial load at zero time point, flies were allowed to rest for 10 min after bacterial injection before plating as described above. The agar plate was maintained at 37°C for 18 hours before CFUs counting. CFUs were log10 transformed.

### Cell culture, transfection and luciferase assay

S2* cells (a gift from Dahua Chen, Institute of Zoology, CAS) were maintained in Drosophila Schneider’s Medium (Invitrogen) supplemented with 10% heat-inactivated fetal bovine serum (Gibco), 100 units/ml of penicillin, and 100 mg/ml of streptomycin at 28°C. Transient transfection of various plasmids, dsRNA was performed with lipofectamine 3000 (Invitrogen), according to the manufacturer’s manual. Luciferase reporter assays were carried out using a dual-luciferase reporter assay system (Promega). Where indicated, cells were treated with PGN (35 μg/μl, 6 h) purified from Erwinia carotovora carotovora 15 (*Ecc15*) referring to previous study(87).

### qRT-PCR

For quantification of mRNA level, about 20 flies carcass/fat body tissue were dissected in sterile PBS buffer on ice at indicated time points post infection, immediately homogenized in 200 μl cold TRIzol with three fly body volumes of ceramic beads (diameter: 0.5 mm), then supplied additional 300 μl TRIzol to reach total 500 μl volume and samples were stored at −80°C.RNA extraction was referred to the manual of commercial kit (Magen, Hipure Total RAN Plus Micro Kit), this kit can effectively remove genomic DNA contamination. cDNA was synthesized by using the kit (abm, 5X All-In-One MasterMix) with total 1μg isolated RNA as template in a 20 μl reaction system. Quantitative real-time PCR (qRT-PCR) was performed using a SYBR green kit (abm, EvaGreen supermaster Mix) on an ABI 7500 or ViiATM 7 thermocycler (Life Technology). Samples from at least four independent biological replicates per genotype were collected and analyzed. House-keeping gene rp49 as the reference gene for data normalization. Primer data for qRT-PCR are provided in Supplementary Table 1.

### Lipid droplet staining and counting

For lipid droplet staining, adult male carcass/fat body tissues were dissected and fixed in 4% fresh prepared paraformaldehyde (PH=7.5) in PBS for 10 min on ice. Tissues were then rinsed twice with PBS (3 min each time), then incubated in PBS containing 1μg/ml of BODIPY 493/503(Invitrogen) dye or 0.5 μg/ml Nile Red (Sigma) for 30 min on ice, DAPI (1μg/μl, final concentration) was added to stain nuclei at last 5 mins of staining process. After staining, tissues were rinsed three times with PBS (3 mins each time), then mounted in mounting medium (Vector, H-1000) for microscopy analysis. To quantify the average lipid droplet size, the average diameter of the three largest lipid droplets per cell was measured generally, with the exception of *plin1* deficiency associated flies, we measured their biggest lipid droplets in one cell (27). 30 fat body cells of each genotype fly randomly selected from eight confocal images were used to analysis the lipid droplet size. To count the size distribution of lipid droplets, the average percentage of the indicated size range of lipid droplets per cell from 30 fat body cells were determined by using the “Analyze Particles” tool embedded in ImageJ software (https://imagej.nih.gov/ij/). To quantify the fluorescence intensity of GFP on the surface of lipid droplets, confocal images acquired from eight fat bodies were measured by ImageJ software.

### Glyceride detection

Glyceride amounts were measured using a TG Quantification Kit (BIOSINO, TG kit). Briefly, for whole body glyceride quantification, groups of 12 one-week old male flies were collected and weighted (about 10 mg) in a 1.5 ml EP tube, then immediately stored at −80°C for subsequent assay. Stored flies were homogenized in 200 μl lysis buffer (10mM KH2PO4, 1mM EDTA, PH=7.4) with three fly body volumes of ceramic beads, and inactivated in water bath at 75°C for 15 min. The inactivated homogenate was homogenized again for 30 s and kept on ice ready for assay. For each glyceride measurement, 3 μl of homogenate was incubated with 250 μl reaction buffer at 37°C for 10 min. After removal debris by centrifugation (2000 rpm, 2 min), 150 μl of clear supernatant was used to perform a colorimetric assay in 96 well plate (Corning® Costar) for absorbance reading at 505 nm. Glyceride level was normalized with fly weight in each homogenate (unit: nmol/mg.fly). For fat body glyceride quantification, 25 fly’s carcass/fat body tissues were dissected and following assay as described above. Glyceride level was normalized with per 25 flies (unit: nmol/25.fly).

### RU486 treatment

RU486 induction was described as before (88). Briefly, A 10 mg/ml stock solution of RU486 (mifepristone; Sigma) was dissolved in DMSO. Appropriate volumes of RU486 stock solution was diluted with water containing 2% ethanol to final concentration of 50 μg/ml.100 μl of the diluted RU486 solution was dipped onto the surface of fresh food in vials (Diameter: 2 cm). The vials were then allowed to dry at room temperature for half day or 4°C for overnight. Flies were transferred to RU486-contained food and raised in 25°C and fresh food was changed every two days.

### NAC Treatment

N-acetyl-L-cysteine (NAC) (Beyotime) fresh solution was prepared by dissolving 0.5 g of NAC powder in 10 ml distilled water, the solution could be aliquoted into 1 ml per EP tube and frozen or stored at −80 °C. 100 μl of NAC solution was dipped onto the surface of fresh food in vials (Diameter: 2 cm). The vials were then allowed to dry at room temperature for half day or 4°C for overnight. Flies were transferred to NAC-contained food and raised in 25°C and fresh food was changed every day.

### ROS detection

We used two methods to detect ROS in fat body, which are *gstD-GFP* reporter flies and dichlorofluorescein diacetate (DCFH-DA) labeling. The oxidative stress reporter construct *gstD-GFP* for evaluating cellular ROS levels has been describe before (58). Briefly, the carcass/fat body of transgenic flies containing a *gstD-GFP* reporter construct were dissected in sterile PBS, fixed in 4% formaldehyde for 10 min on ice, rinsed twice with ice-chilled PBS (3 min each time), then the flaky fat body cells attached to the inner carcass shell were dissected out to mount and confocal image (Vector, H-1000). DCFH-DA (Beyotime, Reactive Oxygen Species Assay Kit) labeling of fresh dissected carcass/fat body tissues was performed according to the manufacturer’s manual, which based on the ROS-dependent oxidation of DCFH-DA to fluorescent molecule 2’-7’dichlorofluorescein (DCF). In brief, the tissues were incubated with PBS containing 20 μM DCFH-DA for 30 min at 30°C, washed with sterile PBS for three times (3 min each) to remove free DCFH-DA that do not uptake by the cell, then the flaky fat body cells attached to the inner carcass shell immediately were dissected out to mount and confocal image (Vector, H-1000). It should be noted that the slices were confocal imaged using the exact same settings for control and experimental groups. The fluorescence intensity is proportional to the ROS levels, fluorescence intensity of GFP or DCF was quantified by using ImageJ software.

### Microscopy and software

LSM700 (Leica) and Olympus FV-1200 confocal laser scanning microscopy were used for imaging. Captured images were analyzed by implemented soft respectively. ImageJ (https://imagej.nih.gov/ij/) was used for analysis of fluorescence intensity and lipid droplets size.

### Statistical analyses

All replicates are showed as the mean ± SD or mean with range. Statistical significance was determined using a paired Student’s t-test for two measurements, one-way ANOVA (Tukey’s HSD) with a multiple t-tests and Multiple t-tests for pairwise comparisons. Kaplan–Meier test for survival curves comparison. All data processing was used with GraphPad Prism 7.0.

#### Sample size choice

The sample size was determined according to the number of data points. Batches of experiment were carried out to ensure repeatability and the use of enough animals for each data point.

#### Randomization

Measures were taken to ensure randomization. Each experimental batch contained more animals than the number of data points, to ensure randomization and the accidental exclusion of animals. In vitro analyses were usually performed on a specimen from animals at each data point to ensure a minimum of three biological replicates.

#### Blinding

Data collection and data analysis were routinely performed by different people to blind potential bias. All measurement data are expressed as mean ± s.d. to maximally show derivations, unless otherwise specified.

## Acknowledgements

We thank Drs. Xun Huang (Institute of Genetics and Developmental Biology, CAS) for providing stocks of *plin1*^38^, *UAS-plin1*-mcherry, *UAS-plin1-RNAi* flies and valuable comments; Zhiwei Liu (Shanghai Ocean University) for providing stock of *UAS-gstD-GFP*; Zhihua Liu (Institute of Biophysics, CAS) for providing *S. typhimurium (SR-11)* strain; Ms. Song-qing Liu (Institute of Biophysics, CAS) for fly food preparation and stock maintenance. We thank Drs. Hui Xiao, Parag Kundu and Philippe Sansonetti (IPS, CAS) and Chengshu Wang (SIPPE, CAS) for comments and manuscript polishment. This work was supported by grants from the Strategic Priority Research Program of the Chinese Academy of Sciences to H.T (XDB29030301), the National Natural Science Foundation of China to L.P (31870887) and J.Y (31670909) and Shanghai Municipal Science and Technology Major Project to L.P (2019SHZDZX02). L.P is a fellow of CAS Youth Innovation Promotion Association (2012083).

## Author’s contributions

Conceptualization, L.W., J.L. and L.P.; Methodology and Validation, L.W., J.L., L.S., K.Y. and L.P.; Formal Analysis, L.W. J.L., J.Y., and L.P.; Investigation, L.W., J.L. and L.P.; Resources, L.P.; Writing-Original Draft, L.W. and L.P.; Writing-Review & Editing, L.P.; Supervision, H.T. and L.P.; Funding Acquisition, J.Y., H.T. and L.P.

## DECLARATION OF INTERESTS

The authors declare no competing interests.

## SUPPLEMENTAL INFORMATION

Supplemental Information includes four figures, and one table.

## Supplementary Figure Legends

**Fig. S1.**
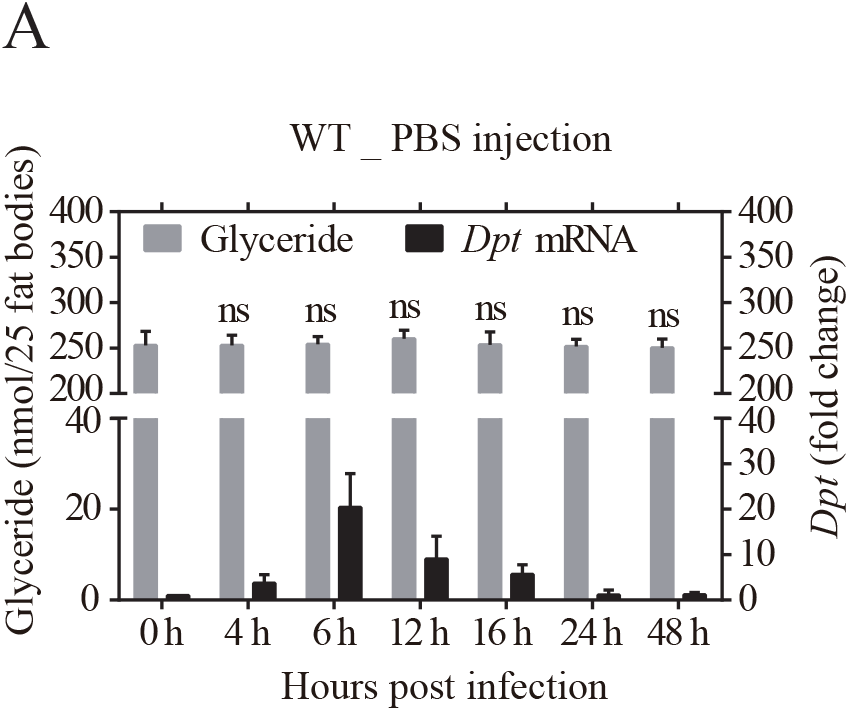
IMD signaling modifies lipid metabolism and LDs morphology. Related to Figure 1. (**A**) Relative *Dptericin* (*Dpt*) mRNA expression (black) and glyceride level (gray) in the fat body of wild type flies at indicated time points after sterile PBS injection. The fold change of mRNA expression was normalized to that of 0 h and four independent repeats (n =20 flies per repeat) were performed at each time point. Total glyceride level of 25 flies’ fat body tissues was quantified in six biological replicates at each time point. Error bar mean ± s.d. Data were analyzed by One-Way ANOVA with Tukey’s multiple-comparison test. \**p* < 0.05; \*\**p* < 0.01; \*\*\**p* < 0.001, ns, no significance.

**Fig. S2.**
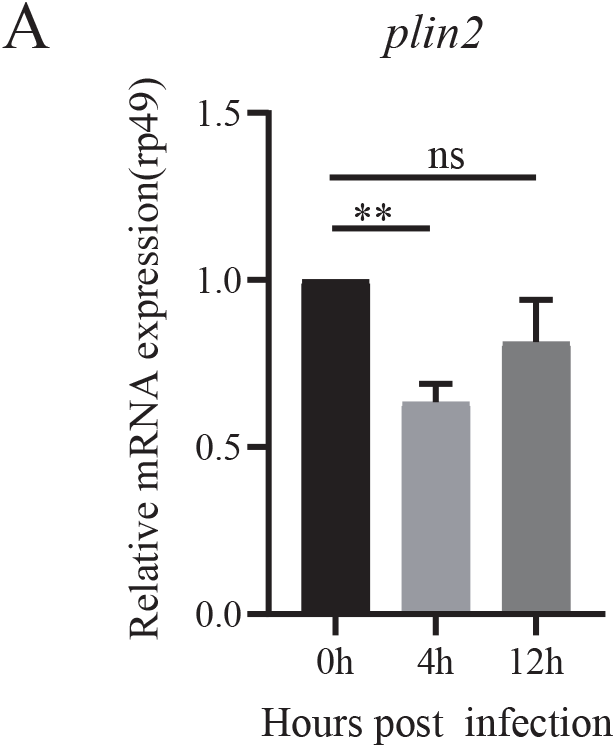
*Plin2* expression of fat body upon *E.coli* infection. Related to Figure 3. (**A**) Relative *plin2* mRNA expression in the fat body of wild type flies at indicated time points post *E.coli* infection. The fold change of mRNA expression was normalized to that of 0 h and at least three independent repeats (n =20 flies per repeat) were performed at each time point. Error bar mean ± s.d. Data were analyzed by One-Way ANOVA with Tukey’s multiple-comparison test. \**p* < 0.05; \*\**p* < 0.01; \*\*\**p* < 0.001, ns, no significance.

**Fig. S3.**
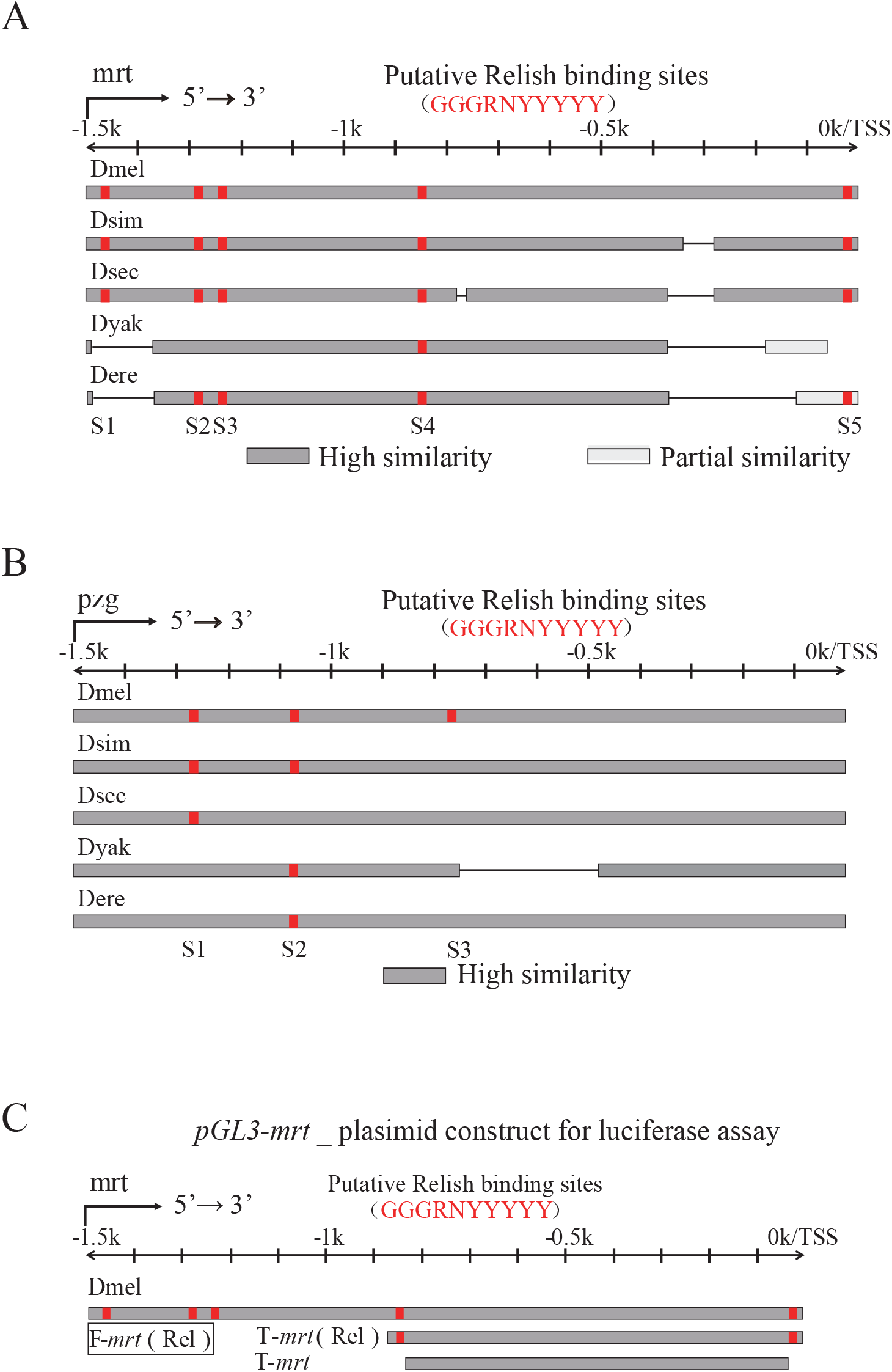
Relish/NF-κB potentially regulates the transcription of *mrt* or *pzg* in *Drosophila* subgroups. Related to Figure 3. (**A and B**) Predicted Relish/NF-κB-binding motifs in the promoter locus of *mrt* (**A**) and *pzg* (**B**) genes of five *Drosophila* subgroups. Dmel, *Drosophila melanogaster*; Dsim, *D. simulans*; sec, *D. sechelia*; Dyak, *D. yakuba*; Dere, *D. erecta*. S1-S5 represent the location site (red color) of conserved binding motifs of Relish. Sequence alignment is analyzed by BLAST in flybase website. TSS: transcription start site. (**C**) Schematic diagram of the *mrt* promoter locus and the plasmid constructs used for luciferase assay. The full length (*mrt*:−1.5k to +1bp), truncated length (T-*mrt* (Rel):−870 to +1bp) and mutant length (T-*mrt*:−870 to +1bp without binding motifs) of *mrt* promoter were indicated.

**Fig. S4.**
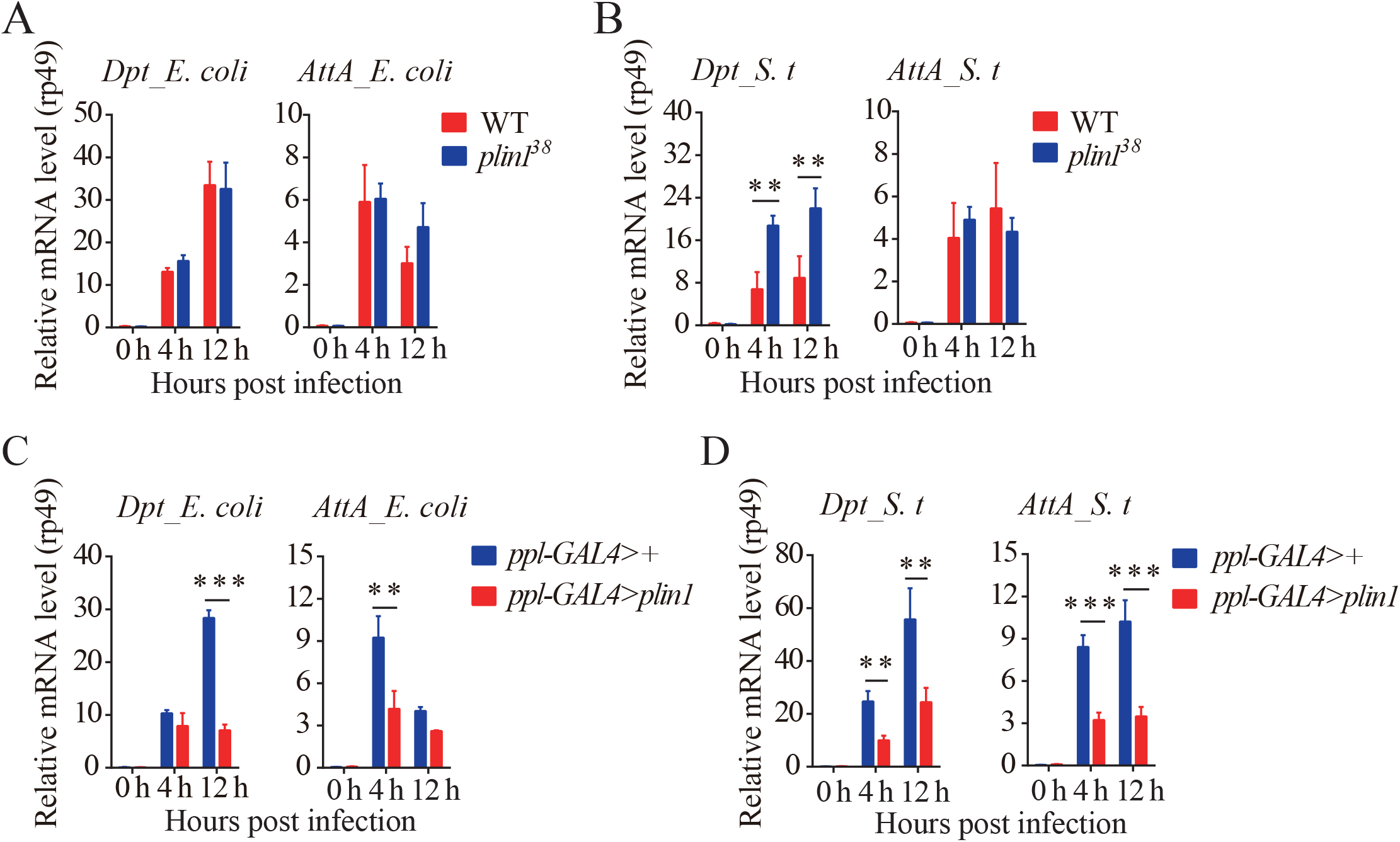
Overexpression *plin1* compromises AMP responses. Related to Figure 4. (**A** and **B**) Relative *diptericin (Dpt)* and *attacin-A (AttA)* mRNA expression in the fat body of *plin1*^38^ mutant flies and wild type flies at indicated time points post *E. coli* (**A**) or *S. typhimurium* (**B**) infection. Four independent repeats were performed (n = 20 per repeat). (**C** and **D**) Relative *diptericin (Dpt)* and *attacin-A (AttA)* mRNA expression of *ppl-GAL4> plin1* flies and *ppl-GAL4> +* control flies at indicated time points post *E. coli* (**C**) and *S. typhimurium* (**D**) infection. Four independent repeats were performed (n = 20 per repeat). Error bars represent mean ± s.d. Data was analyzed by Multiple t-tests. \*\**p* < 0.01; \*\*\**p* < 0.001.

**Table S1:**
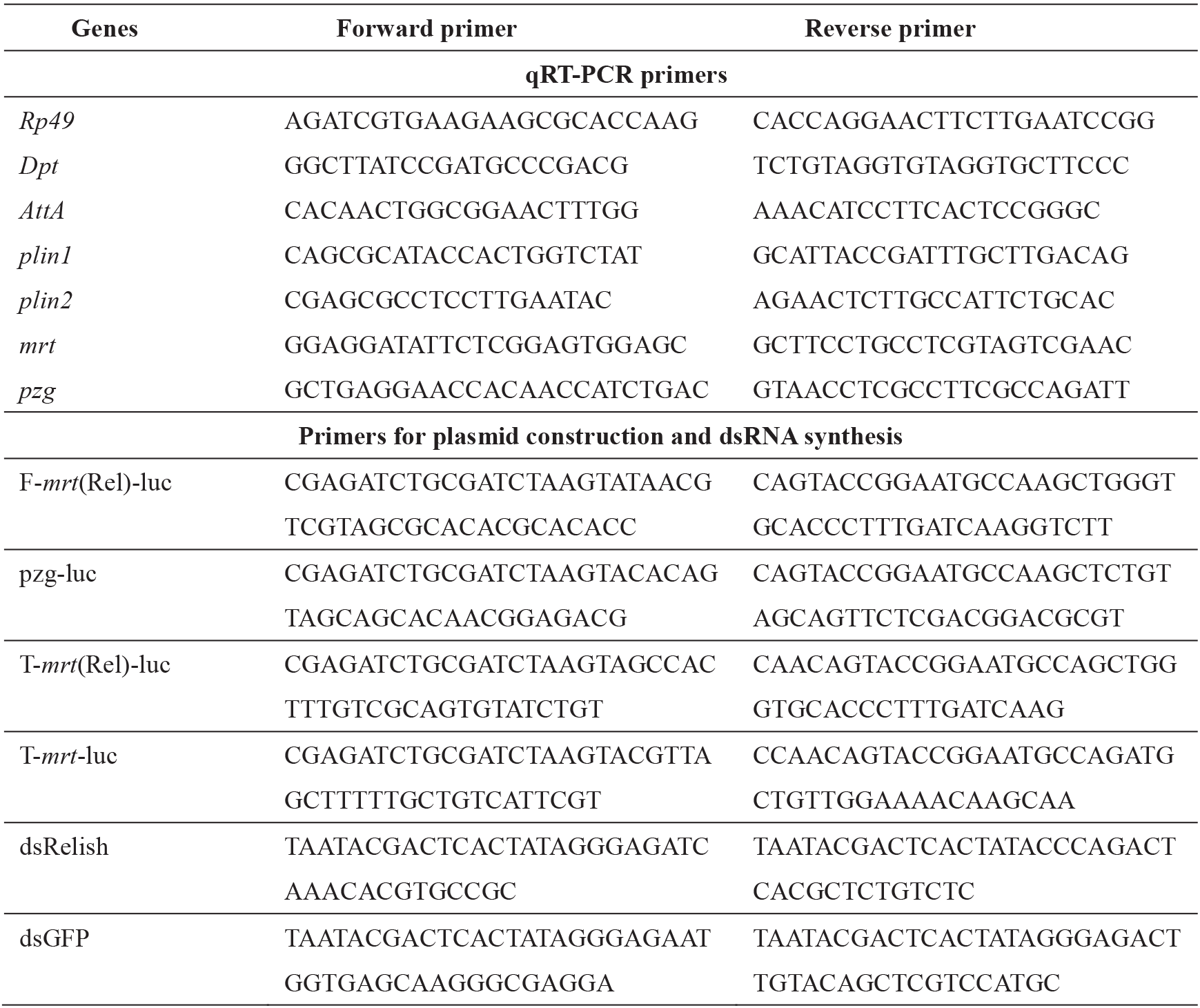
Primers used in this study.

## Notes

### Competing Interest Statement

The authors have declared no competing interest.

